# Quantitative Dual Predation by a Wild-type Bdellovibrio-like Organism and Lytic Bacteriophage Reveals Host-Specific Synergy

**DOI:** 10.1101/2025.11.23.690044

**Authors:** Imelda L. Forteza, Michael Hessari-Bahrami, Bryan John J. Subong

## Abstract

Bacteriophages and wild-type Bdellovibrio-and-like organisms (BALOs) are key microbial predators that influence bacterial population turnover, yet their potential synergy during co-infection remains poorly characterized. Here, we quantified dual predation by a wildtype BALO isolate (*EMS*) and a lytic phage (*M7f*) against *Aeromonas sobria* and *Escherichia coli* using synchronized single-cycle infections. Colony-forming units (CFU), plaque-forming units (PFU), and optical density (OD_600_) were monitored over time, supported by Bliss independence modeling and Kaplan–Meier survival analysis. Dual predation did not accelerate lysis onset but increased the magnitude of prey clearance by approximately 2–3-fold in *E. coli* relative to single-predator treatments. Transmission electron microscopy confirmed co-infection, revealing phage particles adsorbed to prey cells containing BALO-induced bdelloplasts. Together, these findings show that wild-type predators’ complementary infection modes, rapid phage lysis and slower BALO periplasmic invasion, create host-specific synergy. This quantitative framework offers mechanistic insights into natural co-predation and underscores the potential of integrated predator systems for microbial control.

## 1. Introduction

Microorganisms rarely function in isolation. Within shared ecological niches, microbial communities engage in complex interactions—competing or cooperating for survival and access to limited resources [1][2]. Among these interactions, microbial predation plays a central role in shaping community structure, nutrient fluxes, and ecosystem stability [3] [4]. Two key groups of microbial predators, bdellovibrio-and-like organisms (BALOs) and lytic bacteriophages, exemplify distinct yet functionally analogous strategies of bacterial predation [3][5][6][7]. Lytic phages are obligate viruses that hijack host replication machinery and lyse cells from within, whereas BALOs are Gram-negative predatory bacteria that invade the periplasmic space, forming a transient intracellular niche (the bdelloplast) before lysing their host [8] [9]. Despite these mechanistic differences, both predator types drive microbial turnover across terrestrial and aquatic systems [10].

While phage–host dynamics are well characterized, BALOs remain comparatively underexplored. Their ability to traverse biofilms and prey upon a broad range of Gram-negative bacteria highlights their potential as broad-spectrum biocontrol agents [11][10]. However, in natural systems, bacterial populations are often exposed to multiple predators simultaneously, and the ecological consequences of BALO–phage co-predation remain poorly understood [12][13]. Although theoretical models predict additive or synergistic effects, empirical validation under synchronized and controlled conditions is lacking [14].

To address this gap, we systematically evaluated the co-infection dynamics of a novel BALO *EMS* and a lytic phage *M7f* against *Aeromonas sobria* and *Escherichia coli*. Using synchronized, single-cycle infections, we tracked lysis kinetics via colony-forming units (CFU), plaque-forming units (PFU), and optical density (OD_600_). We further employed transmission electron microscopy (TEM) to visualize infection stages, Bliss independence modeling to quantify synergy, and Kaplan–Meier survival analysis to assess infection timing.

Although wild-type Bdellovibrio-and-like organisms (BALOs) and lytic phages frequently co-occur in natural environments, their interaction dynamics remain poorly resolved [15]. Prior work has described dual predation in laboratory strains (e.g., *B. bacteriovorus* HD100 and environmental phage isolates) and rare coinfection events within single prey cells [16] [17] yet no study has quantified co-infection kinetics using synchronized single-cycle assays with wild-type environmental isolates. The objective of this study is to define the mechanistic and quantitative features of dual predation; we focused on time-resolved lysis kinetics rather than genomic characterization. Here, we tested the hypothesis that synergy arises from complementary infection modes: a wild-type BALO that invades and consumes prey periplasmically over several hours [18], and a lytic phage that rapidly infects metabolically active surface-accessible cells [19]. These findings expand our understanding of wild-type microbial predator–prey dynamics and highlight potential applications for combinatorial biocontrol.

## 2. Materials and Methods

### 2.1 Bacterial Strains and Predator Isolation

A bdellovibrio-like organism (BALO), designated *EMS*, was isolated specifically for this study from sewage water in Quezon City, Metro Manila, Philippines, using *Escherichia coli* as prey. The *E. coli* isolate used as the prey organism as well as the *Aeromonas sobria* strain used for propagating phage *M7f* was obtained from previously characterized stock cultures maintained at the University of Santo Tomas Graduate School (UST-GS). The identification of *E. coli* and *A. sobria* was verified using API 20E and API 20NE (bioMérieux) biochemical profiling.

The lytic bacteriophage *M7f* was originally isolated in an earlier study from sewage water in Dumaguete City using *A. sobria* as prey. For the present work, these bacterial isolates and the phage were retrieved from UST-GS stock collections and revived following standard procedures.

All bacterial strains were maintained at ambient temperature (∼25–28 °C) in nutrient broth (NB) supplemented with divalent cations. To purify EMS, mixed cultures with *E. coli* were sequentially filtered through 0.45 µm and 0.2 µm membranes. Single plaques were isolated on double-layer agar (DLA) plates containing 24 mM HEPES and divalent cations, followed by three serial purification passages before storage at 4 °C. Phage *M7f* was purified using the same procedure.

Before experimental use, all UST-GS stock culture isolates (*E. coli, A. sobria*) were verified for purity and identity. Their previously assigned biochemical profiles (API 20E and API 20NE) were confirmed by performing key phenotypic tests, Gram staining, oxidase reaction, and lactose fermentation on MacConkey agar. Observed characteristics matched expected profiles, validating these isolates for use in this study.

### 2.2 Host Range Screening and Spot Assays

Host range screening determined which of the 29 primary bacterial isolates served as common hosts to both bacteriovores. Actively growing host cultures (5 h in NB) were mixed with molten top agar (0.6%) and poured over bottom agar (1.2%) DLA plates. For each of the 28 BALO isolates and the phage *M7f*, 10 µL of crude filtrate was spotted onto triplicate DLA plates.

Positive controls verified the viability of each bacteriovore with its respective host; negative controls substituted sterile water for predator filtrate. A bacterial isolate was considered a common host if both predators produced plaques within 5 days of incubation at room temperature.

### 2.3 Synchronous Single-Cycle Lysis Assays

Modified protocols were used to quantify competitive predation [20][21] on the shared hosts *A. sobria, E. coli*, and *Klebsiella pneumoniae* (Fig. 7). Treatments included host alone (H), BALO + host (Bd + H), phage + host (BΦ + H), and co-predation (Bd + BΦ + H). Synchronous infections were established in 10 mL HEPES buffer (24 mM, supplemented with Ca^2+^ and Mg^2+^) at predator-to-prey ratios of 1:1 to 1:10. Cultures were incubated at 28 °C with shaking (150 rpm). One ml samples were withdrawn at 0, 0.5, 1, 2, 3, 4, and 5 h post-infection for quantification. Each treatment was measured in triplicate.

### 2.4 Quantification of Lysis Kinetics

Lysis dynamics were quantified using three complementary metrics: CFU, PFU, and OD, each measured in triplicate per time point.

#### 2.4.1 Colony-forming units (CFU mL^−1^)

Samples were serially diluted (10^−1^–10^−7^) in sterile buffer and pour-plated on MacConkey agar. Plates were incubated at 37 °C for 24h before counting viable prey cells.

#### 2.4.2 Plaque-forming units (PFU mL^−1^)

BALO and phage titers were determined by DLA assays using *E. coli* as indicator prey. For BALO quantification, samples were filtered through 0.45 µm membranes to remove phages; for phage quantification, through 0.45 µm + 0.2 µm filters to remove BALOs. Plates were incubated at room temperature and read after 1–2 days for phages and 3–4 days for BALOs.

#### 2.4.3 Optical density (OD)

Turbidity was monitored at 600 nm using a Spectronic GENESYS 5 spectrophotometer, with sterile HEPES buffer as a blank. OD trends served as a proxy for bulk lysis dynamics. Derived parameters included burst size (ratio of PFU mL^−1^ increase to CFU mL^−1^ decline during exponential infection), effective multiplicity of infection (eMOI, PFU/CFU ratio), and latent period (interval between infection onset and the first significant PFU rise). Summary statistics are provided in Supplementary Tables S1–S6.

### 2.5 Transmission Electron Microscopy

Transmission electron microscopy (TEM) visualized interactions between BALO *EMS* and phage *M7f* infecting the shared host *A. sobria* (PM2f). Washed pellets of *A. sobria* were resuspended in 0.5 mL HEPES and mixed with equal volumes (0.5 mL each) of host-free pure filtrates of both bacteriovores. After vortex-mixing for 30 min, mixtures were centrifuged at 16 000 rpm for 30 min, washed twice in HEPES, and fixed in 1.5% glutaraldehyde. Samples were post-fixed in osmium tetroxide (OsO_4_), dehydrated in graded acetone, embedded in resin, and polymerized overnight. Ultra-thin sections were stained and viewed at 100 kV on a JEOL JEM-TEM (x) at the Electron Microscope Laboratory, St. Luke’s Medical Center (Manila, Philippines).

### 2.6 Statistical Analysis

All experiments were performed in triplicate and at least three biological replicates, each derived from independent colonies processed in parallel. Statistical analyses were conducted in R (v4.1.1; R Core Team 2021) within RStudio (v2023.09.0) using ggplot2 [22], tidyverse [23], scales [24], survival [25], and survminer [26].

Kaplan–Meier survival curves were used to compare lysis timing between treatments, and significance was assessed via log-rank tests. Pearson’s correlation and simple linear regression were used to quantify OD–CFU and PFU–CFU relationships, reporting r, R^2^, and *p* values (Fig. S1). Regression significance was evaluated with an F-test. Principal Component Analysis (PCA) was performed on the correlation matrix (prcomp function) to visualize multivariate relationships among burst size, lysis efficiency, and synergy index (Fig. 3). Predator interaction strength was quantified using the Bliss independence model to compute a synergy index (Table S2). No formal correction for multiple comparisons was applied, as analyses targeted specific a priori hypotheses.

All R scripts used for statistical analysis and figure generation are openly available on Zenodo under DOI: https://doi.org/10.5281/zenodo.17626134.

## 3. Results

**Table 1.** Preliminary Predator-Prey Screening Matrix.

| Presence of Interaction Across Prey Hosts and Predator Types |  |  |  |
| --- | --- | --- | --- |
| Prey Code | Prey Host | Phage M7f | BALO EMS |
| PM1b | *Klebsiella pneumoniae* |  |  |
| PM1e | *Vibrio mimicus* |  |  |
| PM1F | *Burkholderia cepacia* |  |  |
| PM2a | *Klebsiella pneumoniae* | * | * |
| PM2b | *Klebsiella pneumoniae* |  |  |
| PM2c | *Klebsiella pneumoniae* |  |  |
| PM2d | *Vibrio hollisae* |  | * |
| PM2e | *Burkholderia cepacia* |  |  |
| PM2f | *Enterobacter aerogenes* |  |  |
| PM3a | *Enterobacter cloacae* |  |  |
| PM3b | *Escherichia coli* |  |  |
| PM3c | *Kluyvera sp.* |  |  |
| PM3e | *Alcaligenes xylosoxidans* |  | * |
| PM4c | *Klebsiella pneumoniae* |  |  |
| PM4d | *Klebsiella pneumoniae* |  |  |
| PM6d2 | *Cryseomonas luteola* |  |  |
| PM7f | *Aeromonas sobria* | * | * |
| SH3a | *Escherichia coli* |  |  |
| SH3b | *Escherichia coli* |  |  |
| SH3c | *Stenotrophomonas maltophilia* |  |  |
| EMS | *Escherichia coli* | * | * |
| SH6f | *Escherichia coli* |  |  |
| SH8c | *Chryseomona luteola* |  |  |
| BL3b | *Enterobacter cloacae* |  |  |
| BL3c1 | *Shewanella putrefaciens* |  |  |
| BL3c2 | *Aeromonas sp.* |  |  |
| BL3e | *Enterobacter cloacae* |  |  |
| BL3h | *Enterobacter cloacae* |  |  |
| **Notes:** PM2a: Secondary host of Phage M7f and BALO EMS.<br>PM3e: Secondary host of BALO EMS.<br>EMS: Primary host of BALO and secondary host of Phage M7f.<br>PM2d: Secondary host of BALO. |  |  |  |

### 3.1 Isolation and Screening of Bacteriovores

BALO *EMS* and lytic phage *M7f* were isolated from wastewater and tested against 29 Gram-negative bacterial isolates. Plaque assays showed both predators formed clearings on a shared subset of three prey strains, *Klebsiella pneumoniae, Escherichia coli*, and *Aeromonas sobria*, confirming them as common hosts for follow-up assays.

### 3.2 Host-Specific Lysis Kinetics Reveal Mechanistic Complementarity

We measured the dynamics of predation over a 5-hour period in a synchronous infection scenario (Fig. 1). The CFU, PFU, and OD data showed that host abundance was rapidly declining while predator abundance was concurrently increasing (Table S1). At the same time, the single-cycle tests monitored bulk lysis dynamics, predator replication, and bacterial survival.

**Fig. 1.**
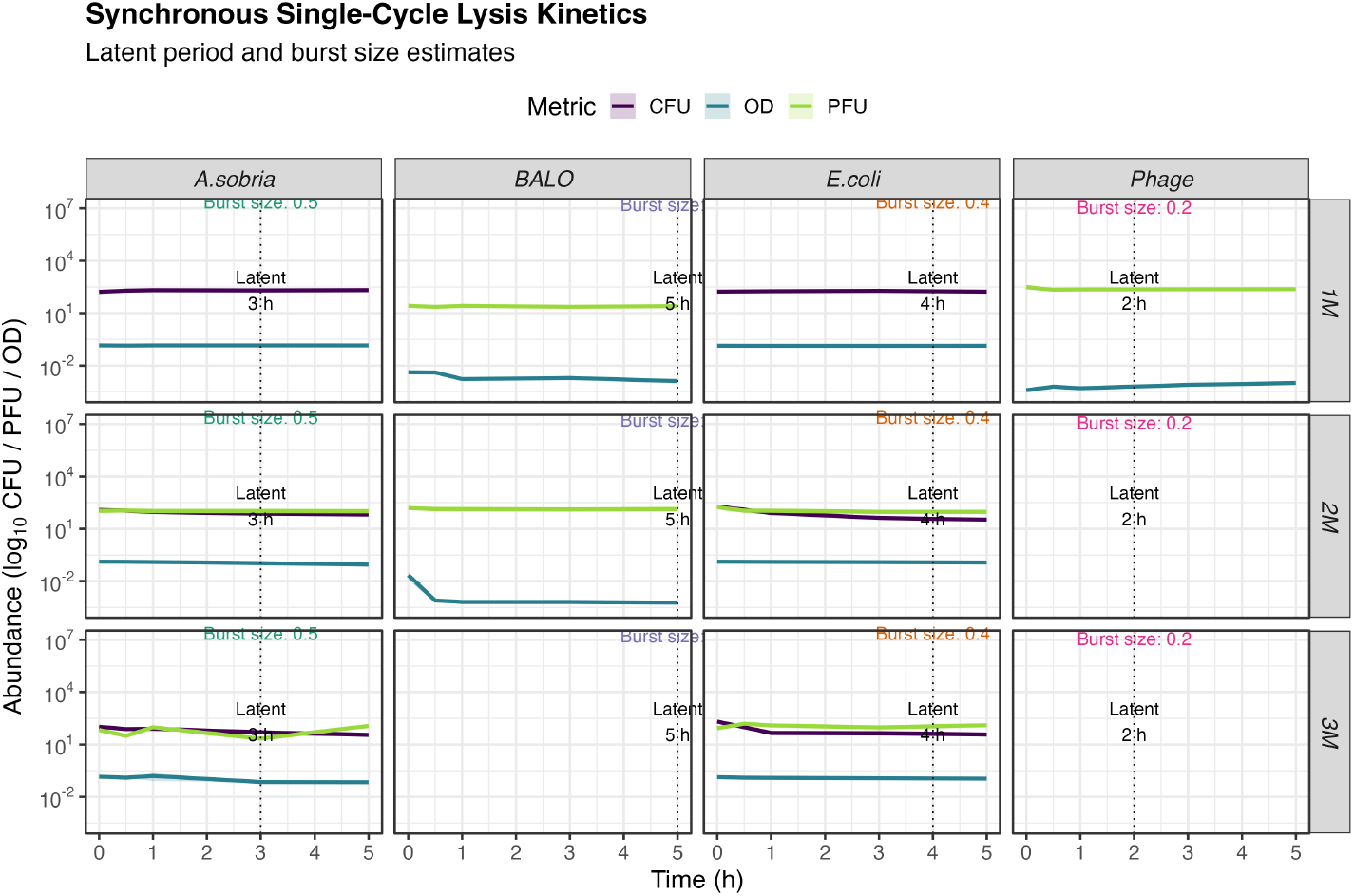
Synchronous single-cycle lysis kinetics of different bacterial hosts under various treatment conditions. Log^10^-transformed CFU mL^−1^ (purple), PFU mL^−1^ (green), and OD_600_ (dashed blue) values are shown with shaded ribbons representing 95% confidence intervals (shaded), plotted over time. The latent period (dotted vertical line) marks the timepoint of first significant PFU mL^−1^ increase. Burst size was estimated as the net PFU mL^−1^ increase per lysed CFU mL^−1^. Each panel represents a specific Host × Treatment combination. OD_600_ provides a bulk view of lysis, while PFU mL^−1^ and CFU mL^−1^ reflect phage propagation and host viability, respectively.

**Fig. 2.**
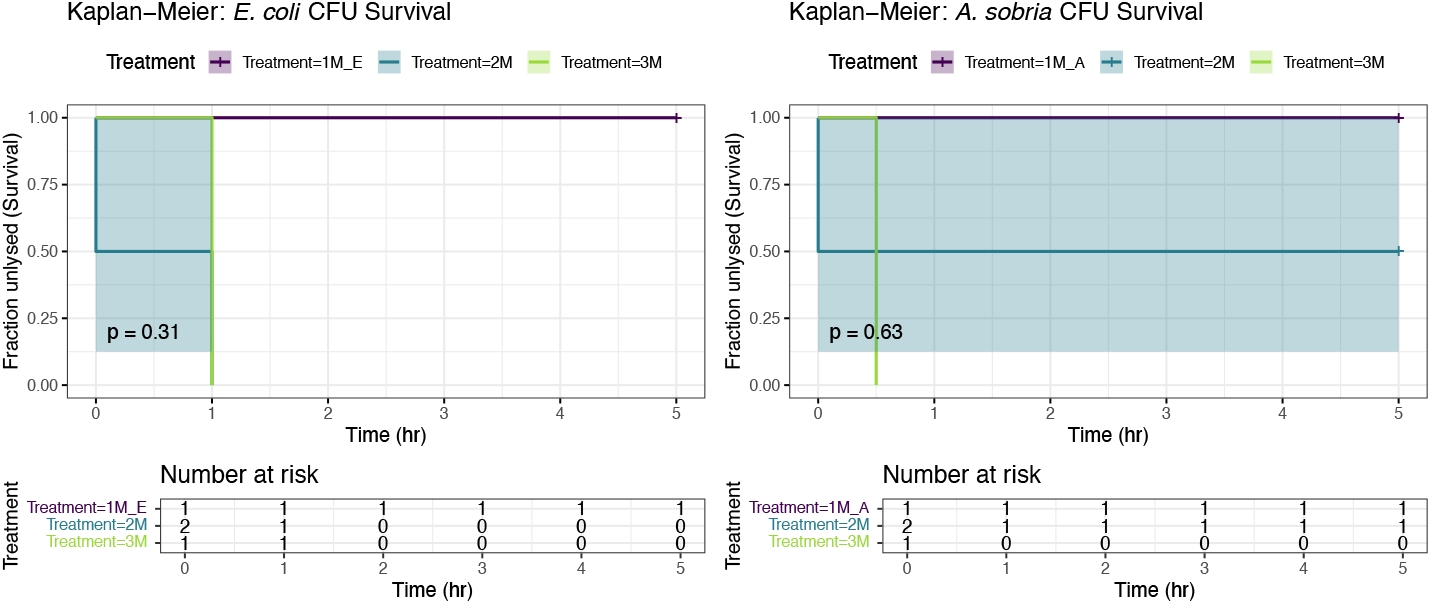
Kaplan-Meier survival curves of *A. sobria* and *E. coli* based on CFU measurements over a 5-hour infection period. Survival is defined as the fraction of unlysed cells at each time point, using CFU mL^−1^ as a proxy for viability. Treatments include 1M (control), 2M, and 3M treatments.

**Fig. 3.**
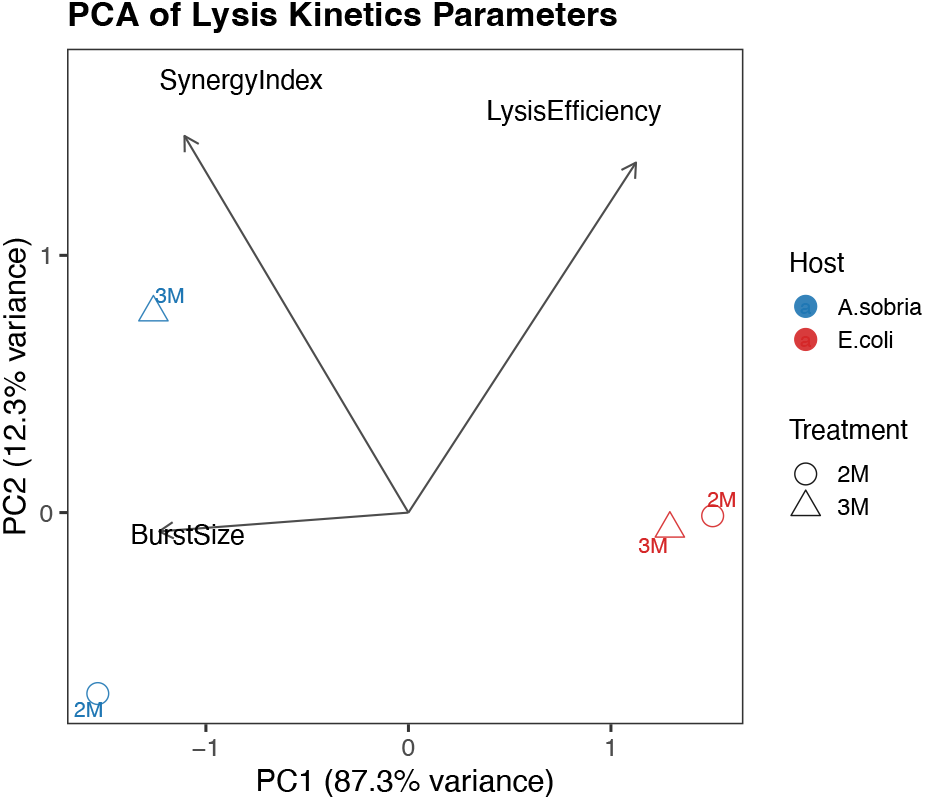
Principal Component Analysis (PCA) of infection kinetics parameters across host– treatment combinations. The first two principal components explain 87.3% and 12.3% of the variance, respectively. *A. sobria* (blue) and *E. coli* (red) are separated along PC1, driven mainly by lysis efficiency and synergy index. Treatments (2M = circle, 3M = triangle) cluster closely within hosts, indicating host-specific rather than treatment differentiation.

**Fig. 4.**
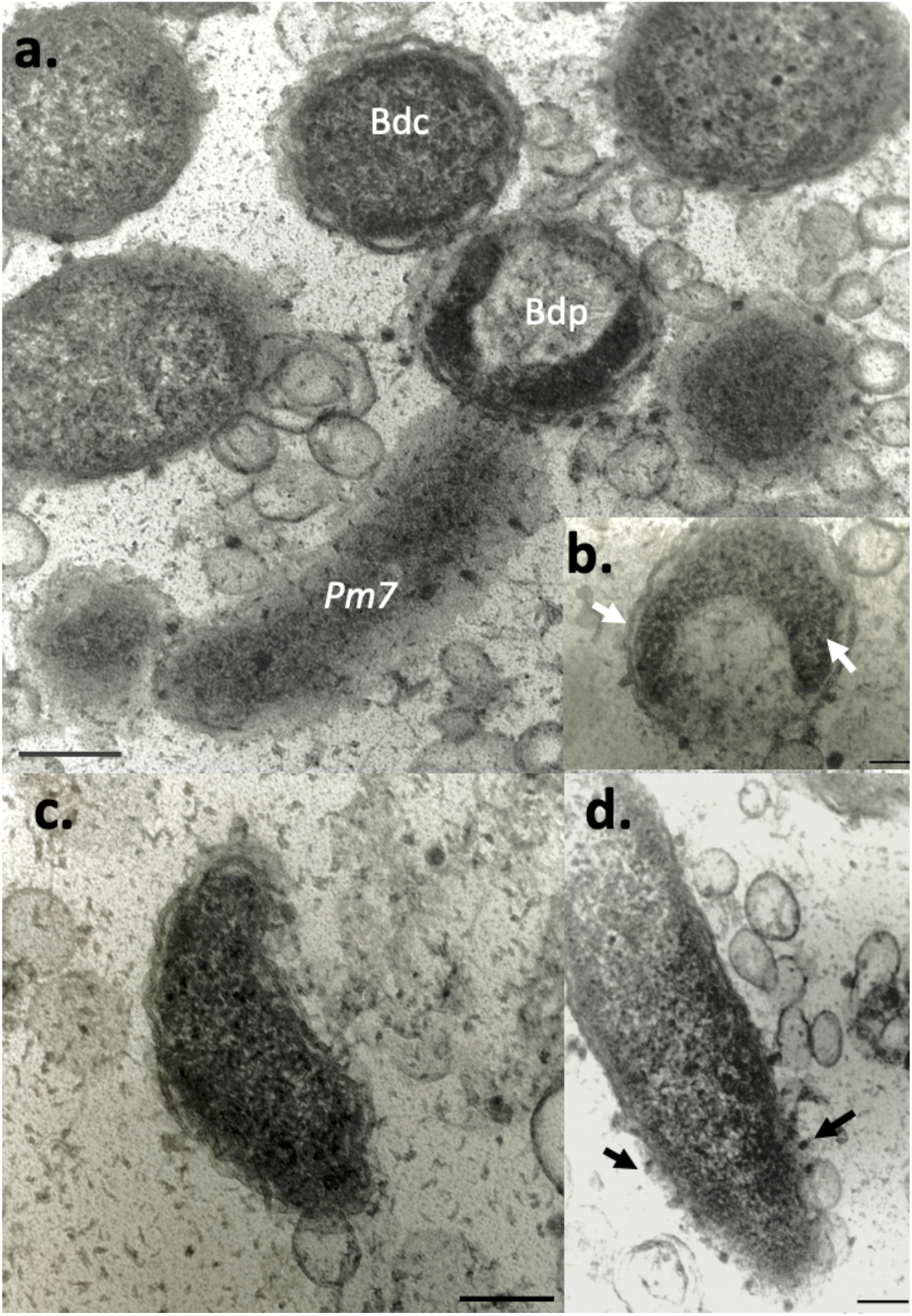
Transmission electron micrographs (TEM) of thin sections showing Bdellovibrio-and-like organism (BALO) *EMS* and lytic bacteriophage *M7f* during co-infection of prey *Aeromonas sobria* (PM7f). (a) Overview of prey cells and infection stages, including bdellocyst (Bdc), bdelloplast (Bdp), and *A. sobria* (PM7f). A lower electron density and crenulate wall distinguish Bdc from Bdp. Scale bar = 200 nm. (b) Bdelloplast structure showing internal organization and external attachment. (c) BALO *EMS* in its comma-shaped attack phase. Magnification ×87,000 (d) *A. sobria* cell with multiple phage *M7f* particles attached to its surface (black arrows). Magnification ×105,000.

Phage infections progressed more quickly, while BALO infections displayed delayed lytic responses (∼5 h). Latent durations were approximately 3 h for *A. sobria* and 4 h for *E. coli*. The estimated burst size varied between 2.07 and 3.07 (Table 2). Figure S4 illustrates effective MOI trajectories.

**Table 2.** Burst size comparison: Plaque counts increase per colony formed between 2M and 3M treatments.

| Burst Size Estimates |  |  |  |  |
| --- | --- | --- | --- | --- |
| Maximum PFU and Burst Size Derived from CFU at Time Zero |  |  |  |  |
| Host | Treatment | Initial CFU (log10) | Max PFU (log10) | Burst Size |
| A.sobria | 2M | 119.39 | 366.00 | 3.07 |
| A.sobria | 3M | 103.11 | 305.00 | 2.96 |
| E.coli | 2M | 187.56 | 389.00 | 2.07 |
| E.coli | 3M | 203.33 | 456.00 | 2.24 |

This table summarizes recalculated burst size based on maximum PFU and initial CFU measurements.

Together, these patterns show that the wild-type BALO *EMS* and phage *M7f* operate on distinct temporal scales. Their complementary lytic strategies create the mechanistic conditions for synergy during co-infection.

### 3.3 Kaplan-Meier Survival Analysis Reveals Differential Lysis Onsets

Kaplan-Meier survival analysis was done to assess host survival or lysis onset timing under different treatments. The Kaplan-Meier curves of CFU reduction over time demonstrated no significant differences in lysis onset between treatments. Survival probability for *A. sobria* gradually decreased (log-rank *p* = 0.63), whereas *E. coli* showed very minor differences (*p* = 0.31, see Fig. 1). Despite synergy in magnitude, the timing of predation was comparable, suggesting co-infection enhances the extent rather than the onset of lysis. Thus, dual predation enhances the magnitude of host clearance, highlighting that synergy stems from magnitude rather than timing.

### 3.4 Multivariate PCA Reveals Distinct Infection Profiles by Host

Principal Component Analysis of burst size, lysis efficiency, and synergy index explained > 99% of variance (PC1 = 87.3%, PC2 = 12.3%). Host identity dominated clustering, with *A. sobria* and *E. coli* separated along PC1 due mainly to synergy index loadings, while treatment effects contributed secondarily along PC2 (Fig. 3, Table S4). These results indicate that host identity, rather than predator treatment, is the dominant driver of infection trajectories.

### 3.5 *E. coli* Exhibits Strong Synergistic Lysis in Co-Treatment

Bliss independence modeling revealed strong synergistic interactions between BALO and phage treatments in *E. coli* under high-dose (3M) conditions, with synergy indices exceeding additive expectations. In contrast, *A. sobria* showed modest or neutral effects, indicating host-specific synergy. Detailed synergy predictions using the Bliss independence model are provided in Table S2, where a strong synergistic interaction was observed between BALO and phage treatments in *E. coli* under 3M conditions. Similarly, Table S3 summarizes correlation and regression relationships between key infection metrics, highlighting a strong inverse relationship between PFU mL^−1^ and CFU mL^−1^ and a positive correlation between MOI and burst size. This demonstrates that synergy is not universal but emerges selectively when prey physiology aligns with the distinct infection modes of the wildtype BALO and its co-occurring phage.

### 3.6 Optical Density Robustly Correlates with Prey Viability

Reductions in OD corresponded strongly with CFU mL^−1^ decline across treatments (r = 0.96 for *E. coli*, r = 0.94 for *A. sobria*), confirming that turbidity loss reflected active lysis rather than nutrient depletion. Regression analyses explained >90% of OD variance by CFU mL^−1^decline (*p* < 0.05). These results confirm that reductions in OD reflect active cell lysis, not spontaneous death, and reinforce the utility of OD measurements as a proxy for viability in short-term predator-prey assays (see Fig. S1B). Thus, OD_600_ provides a reliable proxy for short-term lysis dynamics in synchronized infection assays.

### 3.7 Phage *M7f* Amplification Mirrors Host Decline in *E. coli*

Prey mortality was tightly linked with phage growth, as seen by the inverse correlation between Phage *M7f* PFU mL^−1^ counts and *E. coli* CFU mL^−1^ (F (1, 13) = 19, *p* = 0.0007). Subsequent variation probably represented cumulative phage replication during overlapping cycles (Fig. S1C). This tight coupling confirms that phage-driven mortality closely tracks prey depletion in the presence of the wild-type BALO.

### 3.8 Ultrastructural Evidence of Co-infection in Prey Cells

Transmission electron microscopy (TEM) images confirmed dual predation on *A. sobria*, showing BALO EMS within bdelloplasts and phage *M7f* particles attached to host surfaces, including bdelloplasts membranes. These micrographs suggest spatial and temporal niche overlap between predators. These images suggest that both predators can occupy the same ecological niche within a prey population, possibly targeting different subpopulations or physiological states. These micrographs corroborate that both predators can access overlapping prey niches, consistent with dual predation observed for wild-type BALOs in natural systems.

## 4. Discussion

This study provides a mechanistic, time-resolved view of dual predation by a bdellovibrio-like organism *EMS* and a lytic phage *M7f* against two Gram-negative hosts. Co-predation increased the magnitude of prey clearance without accelerating its onset, as shown by synchronized infections monitored through CFU mL^−1^, PFU mL^−1^, and OD_600_ measurements [17] and [27], the combined effect exceeded the sum of single-predator outcomes, indicating that synergy arises from complementary attack strategies rather than faster infection kinetics[28] [29].

Quantitative analyses support this interpretation. Kaplan–Meier survival curves showed higher overall clearance under co-predation but no shift in lysis timing relative to single infections. Negative PFU mL^−1^–CFU mL^−1^ correlations confirmed phage-driven mortality, while principal-component analysis highlighted burst size, lysis efficiency, and synergy index as key covarying traits explaining host-specific responses. Reduced niche overlap enables coexistence under resource-partitioning models [30][1], consistent with observed Bliss-independence synergy. Context-dependent patterns were evident: synergy occurred in both hosts but varied with predator ratio, *A. sobria* responded optimally at moderate ratios (2M), while *E. coli* exhibited stronger cooperative gains at higher ratios (3M).

Mechanistically, synergy reflects the coexistence of two biophysically distinct infection modes [31][32]. The BALO *EMS* invades the periplasm, forming a bdelloplast that consumes the host internally over several hours, whereas phage *M7f* attaches externally and rapidly lyses cells through hijacked replication. This “dual-front” strategy partitions the prey population, with BALOs targeting less accessible or structurally protected cells and phages infecting metabolically active, surface-exposed ones [25]. When these modes overlap, predators exploit different prey subpopulations, expanding the range of susceptible cells [33][17]. Host physiology further modulates this interaction: *A. sobria* exhibited faster plaque formation but lower synergy, whereas *E. coli* showed slower kinetics yet higher cooperative gains, likely reflecting variation in receptor density, periplasmic thickness, or growth-phase susceptibility [34] [35].

Transmission electron microscopy corroborated this coexistence, revealing phage particles adsorbed to bdelloplast-bearing cells. These observations challenge the assumption that intracellular BALO stages exclude viral co-infection and instead suggest spatiotemporal overlap where phages exploit metabolically active cells while BALOs invade refractory ones, aligning with models of niche complementarity and sequential exploitation [28]. Collectively, the data indicate that synergy emerges from synchronized yet temporally offset predation within shared prey environments[36][37].

Despite its narrow taxonomic focus, this study offers a quantitative basis for comprehending cooperative microbial predation. Our results show that several infection strategies, such as lytic phage cytoplasmic hijacking and bdellovibrio-like bacterial periplasmic invasion, can coexist and improve overall prey clearance through mechanistic complementarity [17]; [38].

In order to clarify the regulatory mechanisms that underlie synergy, this framework can be further developed through the use of molecular methods such as transcriptomics and proteomics, diverse predator–prey combinations, and multi-cycle infections. Such integrative research will be crucial for bringing the theory of microbial interactions into predictive and practical settings, opening the door for the use of integrated predator systems in ecological modeling and microbial control techniques[39][40].

Because both predators in this study are wild-type environmental isolates, the quantitative framework developed here provides an essential foundation for future genomic and transcriptomic work. Similar to insights gained from *B. bacteriovorus* systems [17] and natural coinfection events [16], pairing our mechanistic data with genome-level characterization of *EMS, M7f*, and their prey would clarify the evolutionary and ecological basis of BALO–phage coexistence. This study therefore contributes to the quantitative baseline required for interpreting future genomic investigations of dual predation by wild-type microbial predators.

## 5. Conclusion

In this work, a replicable framework for investigating interactions between a lytic phage and a Bdellovibrio-like bacteria during dual predation is presented. Without conclusive proof of a hastened initiation of lysis, a combination of kinetic and statistical analysis shows host-specific patterns of improved prey clearance. It appears that the two predators’ distinct infection modes are mutually beneficial, as they facilitate a more consistent reduction in prey [41]. These findings contribute to our understanding of cooperative microbial predation and indicate that mixed-predator systems could be effective models for researching microbial relationships. Further research using extended infection cycles and molecular techniques could shed light on the mechanisms underlying these preliminary findings.

## Author Contributions

I.L.F., Conceptualization, investigation, writing-original draft, statistical data analysis, curation and editing. M.H.B.-writing-original draft and editing. B.J.J.S., investigation, writing-original draft and editing

## Funding

This research received no external funding.

## Acknowledgements

We thank Irineo Dogma Jr, PhD, for his supervision and Ms Evelyn S. Fajardo, RMT, for sharing her expertise, providing the lytic bacteriophage M7f and its host isolate, contributing additional bacterial isolates for screening, and offering valuable advice throughout this project. We also gratefully acknowledge Mr. Rodolfo Forteza for his personal financial support that made this work possible.

## Conflicts of Interest

The authors declare no conflicts of interest.

## Notes

https://doi.org/10.5281/zenodo.17626134

### Competing Interest Statement

The authors have declared no competing interest.

