## Supplementary for "Quantitative Dual Predation by a Wild-type Bdellovibrio-like Organism and Lytic Bacteriophage Reveals Host-Specific Synergy"

Table S1| Summary statistics for kinetics assays across treatments and hosts.

**Table S1. Kinetics Summary Statistics**

Mean, Standard Deviation, and Replicates for CFU, PFU, and OD Measurements

| Metric | Time (hr) | Treatmei | Host | Mean | SD | N |
|---|---|---|---|---|---|---|
| CFU | 0.0 | 1M | A.sobria | 165.00 | 78.49 | 9 |
| CFU | 0.0 | 1M | E.coli | 170.78 | 45.95 | 9 |
| CFU | 0.0 | 2M | A.sobria | 119.39 | 50.47 | 18 |
| CFU | 0.0 | 2M | E.coli | 187.56 | 46.42 | 18 |
| CFU | 0.0 | 3M | A.sobria | 103.11 | 44.40 | 9 |
| CFU | 0.0 | 3M | E.coli | 203.33 | 29.40 | 9 |
| CFU | 0.5 | 1M | A.sobria | 197.56 | 71.61 | 9 |
| CFU | 0.5 | 2M | A.sobria | 110.89 | 36.68 | 18 |
| CFU | 0.5 | 3M | A.sobria | 76.11 | 28.30 | 9 |
| CFU | 1.0 | 1M | A.sobria | 208.00 | 70.56 | 9 |
| CFU | 1.0 | 1M | E.coli | 181.44 | 29.55 | 9 |
| CFU | 1.0 | 2M | A.sobria | 92.11 | 31.90 | 18 |
| CFU | 1.0 | 2M | E.coli | 81.39 | 43.35 | 18 |
| CFU | 1.0 | 3M | A.sobria | 79.44 | 20.19 | 9 |
| CFU | 1.0 | 3M | E.coli | 46.11 | 9.52 | 9 |
| CFU | 3.0 | 1M | A.sobria | 203.11 | 52.65 | 9 |
| CFU | 3.0 | 1M | E.coli | 193.11 | 32.13 | 9 |
| CFU | 3.0 | 2M | A.sobria | 73.28 | 26.49 | 18 |
| CFU | 3.0 | 2M | E.coli | 42.22 | 11.58 | 18 |
| CFU | 3.0 | 3M | A.sobria | 49.78 | 13.24 | 9 |
| CFU | 3.0 | 3M | E.coli | 44.33 | 13.95 | 9 |
| CFU | 5.0 | 1M | A.sobria | 209.67 | 59.14 | 9 |
| CFU | 5.0 | 1M | E.coli | 169.56 | 60.89 | 9 |
| CFU | 5.0 | 2M | A.sobria | 66.33 | 36.55 | 18 |
| CFU | 5.0 | 2M | E.coli | 33.44 | 6.43 | 18 |
| CFU | 5.0 | 3M | A.sobria | 35.89 | 10.12 | 9 |
| CFU | 5.0 | 3M | E.coli | 37.33 | 12.41 | 9 |
| OD | 0.0 | 1M | A.sobria | 0.14 | 0.00 | 9 |
| OD | 0.0 | 1M | BALO | 0.00 | 0.00 | 18 |
| OD | 0.0 | 1M | E.coli | 0.13 | 0.00 | 9 |
| OD | 0.0 | 1M | Phage | 0.00 | 0.00 | 18 |
| OD | 0.0 | 2M | A.sobria | 0.13 | 0.03 | 18 |
| OD | 0.0 | 2M | BALO | 0.02 | 0.05 | 20 |
| OD | 0.0 | 2M | E.coli | 0.13 | 0.00 | 18 |
| OD | 0.0 | 3M | A.sobria | 0.14 | 0.00 | 9 |
| OD | 0.0 | 3M | E.coli | 0.13 | 0.00 | 9 |
| OD | 0.5 | 1M | A.sobria | 0.14 | 0.00 | 9 |

Mean, standard deviation (SD), and replicate number (N) are shown for CFU, PFU, and OD measurements over the 5-hour lysis experiment. Data include multiple treatment conditions (1M: single predator; 2M: *Bdellovibrio*; 3M: dual predator) across *A. sobria* and *E. coli* hosts. The table supports kinetic analyses shown in main figures by quantifying variability and replication across time points and metrics.

Table S2| Statistical comparison of treatments versus 1M controls at 0.5 hours.

**Table S2. Treatment Comparison Statistics at 0.5 hr**

**Comparison of treatments vs. 1M controls for CFU, PFU, and OD measurements**

| Host | Treatme | Metric | F-statistic | p-value | Mean (0.5 hr) | Control | Control Mean | Diff |
|---|---|---|---|---|---|---|---|---|
| A.sobria | 2M | CFU | 97.20 | 3.3e-29 | 92.40 | 1M_A | 196.67 | |
| A.sobria | 2M | OD | 3.23 | 4.2e-02 | 0.12 | 1M_A | 0.14 | |
| A.sobria | 2M | PFU | 4.84 | 2.9e-02 | 105.93 | NA | NA | |
| A.sobria | 3M | CFU | 97.20 | 3.3e-29 | 68.87 | 1M_A | 196.67 | |
| A.sobria | 3M | OD | 3.23 | 4.2e-02 | 0.11 | 1M_A | 0.14 | |
| A.sobria | 3M | PFU | 4.84 | 2.9e-02 | 75.84 | NA | NA | |
| BALO | 1M | OD | NA | NA | 0.00 | 1M_BALO | 0.00 | |
| BALO | 2M | PFU | 79.85 | 1.4e-14 | 148.74 | 1M_BALO | 24.90 | [illegible] |
| BALO | 2M | OD | NA | NA | 0.00 | NA | NA | |
| BALO | 2M | PFU | NA | NA | 136.34 | NA | NA | |
| E.coli | 2M | CFU | 28.26 | 4.8e-11 | 86.15 | 1M_E | 178.72 | |
| E.coli | 2M | OD | 35.38 | 1.2e-13 | 0.13 | 1M_E | 0.13 | |
| E.coli | 2M | PFU | 0.01 | 9.2e-01 | 114.99 | NA | NA | |
| E.coli | 3M | CFU | 28.26 | 4.8e-11 | 82.78 | 1M_E | 178.72 | |
| E.coli | 3M | OD | 35.38 | 1.2e-13 | 0.12 | 1M_E | 0.13 | |
| E.coli | 3M | PFU | 0.01 | 9.2e-01 | 116.83 | NA | NA | |

Summary of F-statistics, p-values, and group means for CFU, PFU, and OD measurements across hosts (*A. sobria*, *E. coli*) and treatments (2M, 3M), benchmarked against 1M single predator controls. Significant reductions in CFU and OD, particularly in *A. sobria* and *E. coli* under 2M and 3M, highlight early lytic activity. PFU comparisons show elevated viral levels in treated groups, with statistical significance observed in most comparisons ($p < 0.05$).

Table S3| Descriptive statistics for CFU, PFU, and OD metrics at 0.5 and 5 hours post-infection.

**Table S3. Kinetics Summary at 0.5 hr and 5 hr**

**Descriptive statistics for CFU, PFU, and OD metrics across time points**

| Host | Treatme[illegible] | Metric | Mean (0.5 hr) | SD (0.5 hr) | CV (%) (0.5 hr) | Mean (0 hr) | Mean (5 hr) | % Cha[illegible] (0 to [illegible] |
|---|---|---|---|---|---|---|---|---|
| A.sobria | 1M | CFU | 196.67 | 66.12 | 33.62 | 165.00 | 209.67 | 2[illegible] |
| A.sobria | 1M | OD | 0.14 | 0.00 | 1.83 | 0.14 | 0.14 | -[illegible] |
| A.sobria | 2M | CFU | 92.40 | 41.89 | 45.34 | 119.39 | 66.33 | -4[illegible] |
| A.sobria | 2M | OD | 0.12 | 0.02 | 19.69 | 0.13 | 0.09 | -3[illegible] |
| A.sobria | 2M | PFU | 105.93 | 104.49 | 98.63 | 108.06 | 101.44 | -[illegible] |
| A.sobria | 3M | CFU | 68.87 | 34.63 | 50.28 | 103.11 | 35.89 | -6[illegible] |
| A.sobria | 3M | OD | 0.11 | 0.11 | 95.31 | 0.14 | 0.07 | -5[illegible] |
| A.sobria | 3M | PFU | 75.84 | 76.95 | 101.45 | 67.78 | 115.11 | 6[illegible] |
| BALO | 1M | OD | 0.00 | 0.00 | 70.15 | 0.00 | 0.00 | -6[illegible] |
| BALO | 1M | PFU | 24.90 | 6.72 | 27.00 | 26.72 | 25.56 | -[illegible] |
| BALO | 2M | PFU | 148.74 | 132.24 | 88.91 | 148.74 | NA | |
| BALO | 2M | OD | 0.01 | 0.02 | 471.93 | 0.02 | 0.00 | -9[illegible] |
| BALO | 2M | PFU | 136.59 | 129.11 | 94.53 | 160.16 | 135.16 | -1[illegible] |
| BALO | 2M | OD | 0.00 | NA | NA | NA | NA | |
| BALO | 2M | PFU | 134.32 | 120.25 | 89.53 | NA | NA | |
| E.coli | 1M | CFU | 178.72 | 43.11 | 24.12 | 170.78 | 169.56 | -[illegible] |
| E.coli | 1M | OD | 0.13 | 0.00 | 1.25 | 0.13 | 0.13 | -[illegible] |
| E.coli | 2M | CFU | 86.15 | 69.38 | 80.54 | 187.56 | 33.44 | -8[illegible] |
| E.coli | 2M | OD | 0.13 | 0.01 | 4.86 | 0.13 | 0.12 | -1[illegible] |
| E.coli | 2M | PFU | 114.99 | 118.18 | 102.77 | 170.72 | 93.17 | -4[illegible] |
| E.coli | 3M | CFU | 82.78 | 72.75 | 87.88 | 203.33 | 37.33 | -8[illegible] |
| E.coli | 3M | OD | 0.12 | 0.01 | 6.38 | 0.13 | 0.11 | -1[illegible] |
| E.coli | 3M | PFU | 116.83 | 134.49 | 115.11 | 85.06 | 127.00 | 4[illegible] |
| Phage | 1M | OD | 0.00 | 0.00 | 86.05 | 0.00 | 0.00 | 15[illegible] |
| Phage | 1M | PFU | 246.34 | 85.76 | 34.81 | 308.11 | 240.17 | -2[illegible] |

Mean values, standard deviations (SD), and coefficients of variation (CV) are shown for each treatment condition across *A. sobria*, *E. coli*, BALO, and phage-only groups. Percent change from baseline (0 hr to 5 hr) is included to highlight key trends in predator-prey dynamics. Notable reductions in CFU and OD were observed for both host species under 2M and 3M treatments, while PFU values showed substantial increases in dual predator conditions. These kinetics data inform burst size calculations, effective MOI shifts, and synergy estimates in related analyses.

Table S4| Predation summary metrics for dual predator assays.

**Table S4. Predation Summary Metrics**

**Summary of CFU, PFU, Burst Size, Lysis Efficiency, and Synergy Indices used for PCA**

| Host | Treatme | Initial CFU | Max PFU | Burst Size | Lysis Efficiency (%) | Synergy Index |
|---|---|---|---|---|---|---|
| A.sobria | 2M | 119.39 | 366 | 3.07 | 44.44 | 105.91 |
| A.sobria | 3M | 103.11 | 305 | 2.96 | 65.19 | 126.66 |
| E.coli | 2M | 187.56 | 389 | 2.07 | 82.17 | 80.74 |
| E.coli | 3M | 203.33 | 456 | 2.24 | 81.64 | 80.21 |

Overview of initial CFU, peak PFU, estimated burst size, lysis efficiency, and Bliss synergy index for *A. sobria* and *E. coli* under 2M (Bdellovibrio-only) and 3M (dual predator) treatments. Burst size reflects the ratio of PFU increase per unit CFU loss over the lysis period. Lysis efficiency represents the percent reduction in CFU relative to the starting value. Notably, *A. sobria* under 3M treatment showed the highest synergy index (126.66), suggesting enhanced combined predatory effects, whereas *E. coli* exhibited high lysis efficiency but near-additive interaction outcomes.

Table S5| Table Summary of Effective multiplicity of infection (MOI) over time during dual predator lysis assays

**Effective MOI Time Series Data**

CFU, PFU, and Calculated Effective MOI Across Time Points

| Time (hr) | Host | Treatment | CFU (log10) | PFU (log10) | Effective MOI |
|---|---|---|---|---|---|
| 0.00 | **A.sobria** | 2M | 119.39 | 108.06 | 0.91 |
| 0.00 | **A.sobria** | 3M | 103.11 | 67.78 | 0.66 |
| 0.00 | **E.coli** | 2M | 187.56 | 170.72 | 0.91 |
| 0.00 | **E.coli** | 3M | 203.33 | 85.06 | 0.42 |
| 0.50 | **A.sobria** | 2M | 110.89 | 112.17 | 1.01 |
| 0.50 | **A.sobria** | 3M | 76.11 | 32.00 | 0.42 |
| 1.00 | **A.sobria** | 2M | 92.11 | 104.00 | 1.13 |
| 1.00 | **A.sobria** | 3M | 79.44 | 95.78 | 1.21 |
| 1.00 | **E.coli** | 2M | 81.39 | 108.83 | 1.34 |
| 1.00 | **E.coli** | 3M | 46.11 | 123.61 | 2.68 |
| 3.00 | **A.sobria** | 2M | 73.28 | 104.00 | 1.42 |
| 3.00 | **A.sobria** | 3M | 49.78 | 22.00 | 0.44 |
| 3.00 | **E.coli** | 2M | 42.22 | 92.83 | 2.20 |
| 3.00 | **E.coli** | 3M | 44.33 | 94.50 | 2.13 |
| 5.00 | **A.sobria** | 2M | 66.33 | 101.44 | 1.53 |
| 5.00 | **A.sobria** | 3M | 35.89 | 115.11 | 3.21 |
| 5.00 | **E.coli** | 2M | 33.44 | 93.17 | 2.79 |
| 5.00 | **E.coli** | 3M | 37.33 | 127.00 | 3.40 |

This table summarizes bacterial CFU, viral PFU, and calculated effective MOI over the lysis time course for velvet worm gut isolates.

Time-resolved measurements of bacterial density (CFU, $\log_{10}$), viral abundance (PFU, $\log_{10}$), and effective MOI (log PFU/CFU) are shown across a 5-hour infection course for *A. sobria* and *E. coli* under Bdellovibrio-only (2M) and dual predator (3M) treatments. Values reflect dynamics of predator-to-prey ratios, with notable increases in effective MOI over time, particularly in *E. coli* under 3M, indicating elevated lytic pressure.

**Table S6 | Table Summary of Lysis Efficiency with Bliss Energy Indices.**

| Bliss Interaction Analysis<br>Observed Lysis Efficiency and Synergy Indices | | | | |
|---|---|---|---|---|
| Host | Treatment | Observed Lysis Efficiency (%) | Expected Lysis (%) | Bliss Synergy Index |
| **A.sobria** | 2M | 44.44 | −61.47 | 105.91 |
| **A.sobria** | 3M | 65.19 | −61.47 | 126.66 |
| **E.coli** | 2M | 82.17 | 1.43 | 80.74 |
| **E.coli** | 3M | 81.64 | 1.43 | 80.21 |

This table summarizes observed and expected lysis efficiencies along with Bliss synergy indices for combined treatments.

Bliss synergy indices for combined predator treatments across bacterial hosts. Synergy index values were calculated based on observed and expected lysis efficiencies under 2M (Bdellovibrio-only) and 3M (Bdellovibrio + phage) treatments for *A. sobria* and *E. coli*. A Bliss index >100 indicates synergistic interaction, 100 indicates additivity, and <100 suggests antagonism. While E. coli treatments exhibited near-additive or slightly antagonistic effects (Bliss indices ~80), *A. sobria* showed strong synergy under both treatments, with the highest index (126.7) observed in the 3M condition.

Supplementary Figures

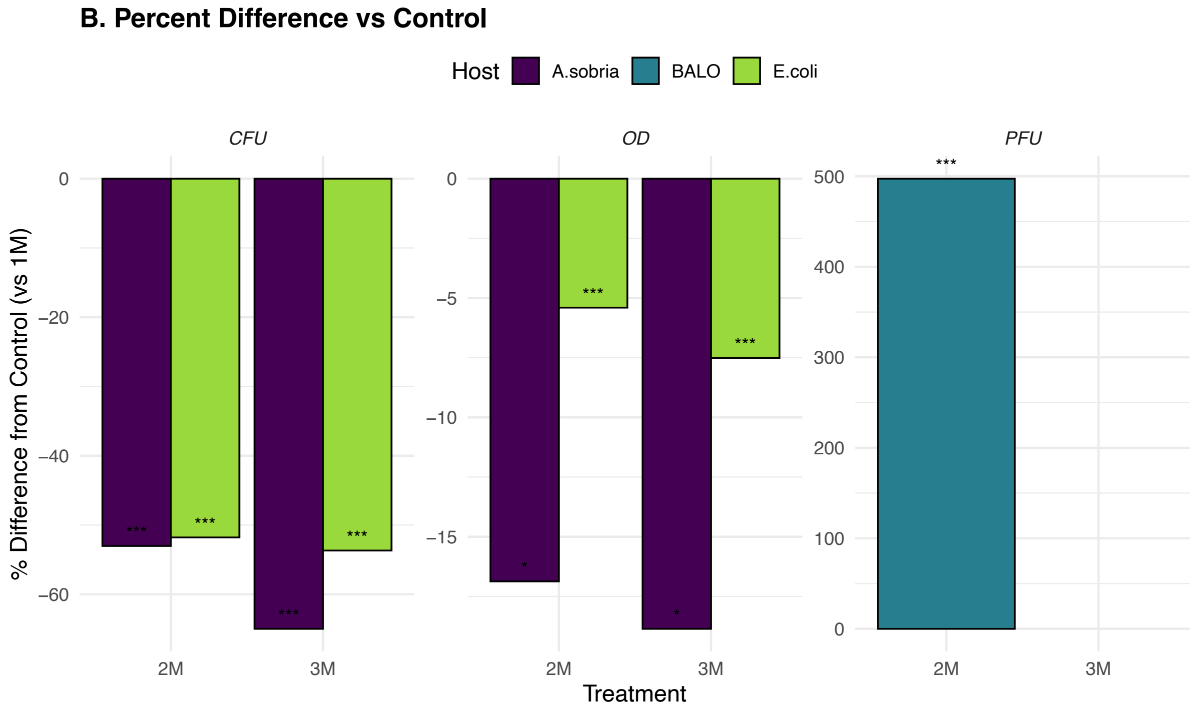


**Fig S1.** *Regression and percent change analyses linking OD, CFU, and PFU dynamics across treatments.*

(A) Percent difference from 1M control values in CFU, OD, and PFU at 5 hours post-infection under 2M and 3M treatments. Bars represent changes relative to single-predator baselines for *A. sobria*, *E. coli*, and BALO. Asterisks denote significance levels ($p < 0.05$, **p** < 0.01, ***p*** < 0.001). Both *A. sobria* and *E. coli* showed substantial CFU reductions, while OD decreases were smaller but still significant. PFU in BALO increased sharply under 2M (~500%), reflecting enhanced lytic activity.
(B) Positive correlations between $OD_{580}$ and CFU in *E. coli* exposed to BALO EMS ($r = 0.96$) and *A. sobria* under dual predator treatment ($r = 0.94$) confirm OD as a reliable proxy for cell lysis.
(C) Phage M7f titers were inversely correlated with *E. coli* CFU over the 5-hour infection course ($R^2 = 0.60$, $p < 0.001$), indicating that phage amplification closely tracks host decline.
Together, these results support the use of OD and PFU as effective indicators of lytic activity and reinforce interpretations in Table S5 (effective MOI time series) and Table 2 (burst size estimates).

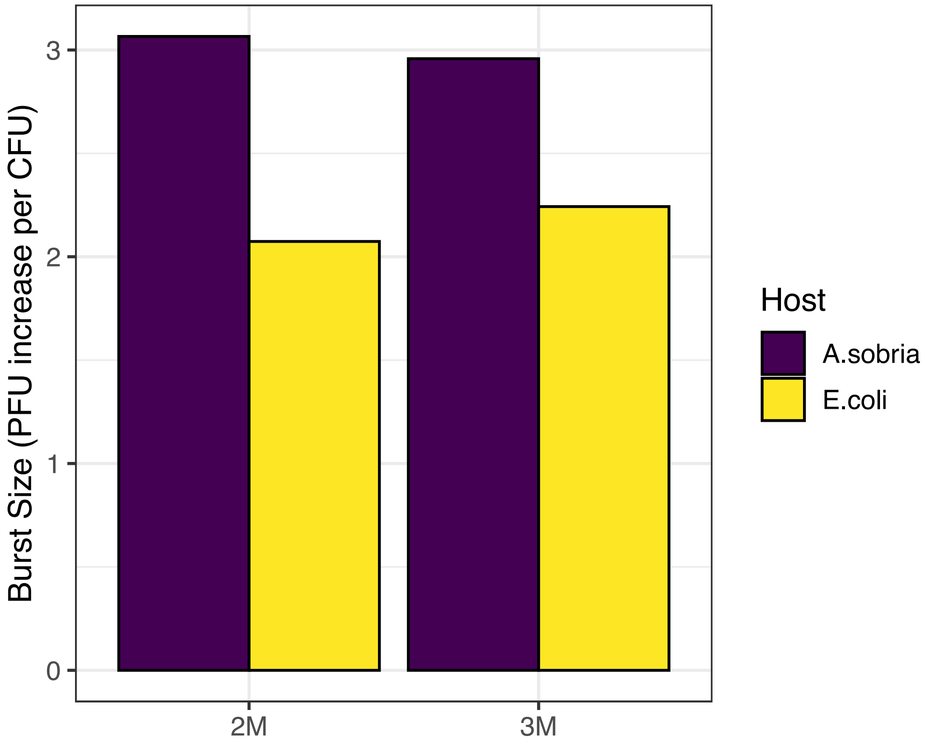


Fig. S2 *Estimated burst size of dual predator systems across Gram-negative hosts.* Burst size was calculated as the increase in PFU per unit decrease in CFU over the 5-hour infection period. Bars represent mean values for each treatment group, with individual comparisons shown for *A. sobria* and *E. coli*. Results highlight host-specific differences in phage and BALO-induced lytic productivity.

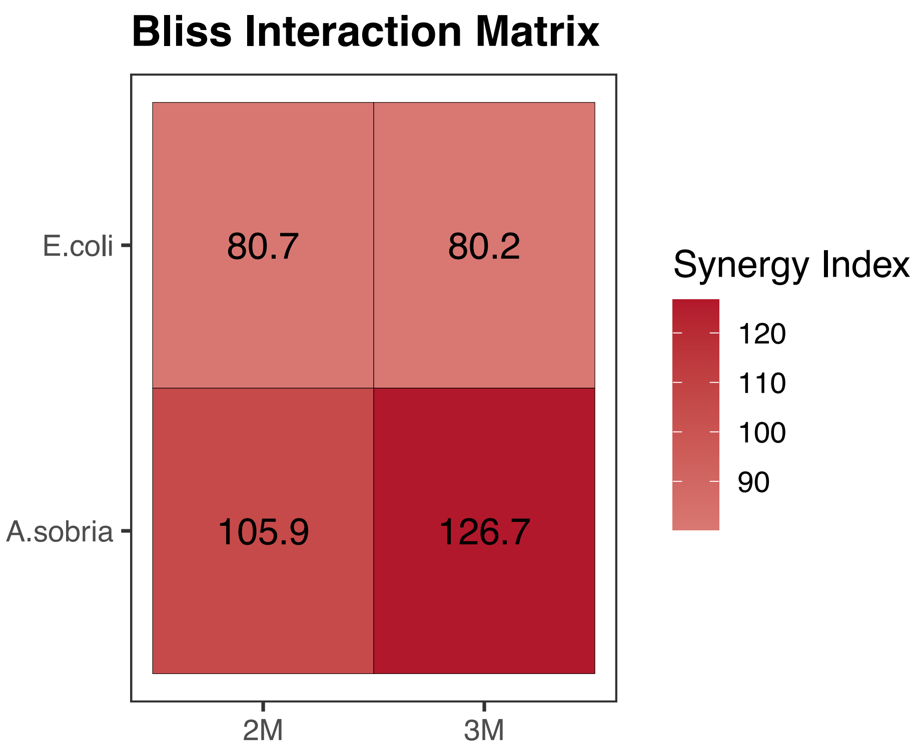


Fig. S3 | Bliss independence synergy index for dual predator treatments across hosts. Heatmap shows synergy index values for *E. coli* and *A. sobria* under 2M (Bdellovibrio-only) and 3M (Bdellovibrio + phage) treatments. A Bliss index >100 indicates synergy, 100 indicates additive effects, and <100 suggests antagonism. Strongest synergistic interaction (Bliss index = 126.7) was observed in *A. sobria* under the 3M treatment, suggesting enhanced dual predator efficacy in this host.

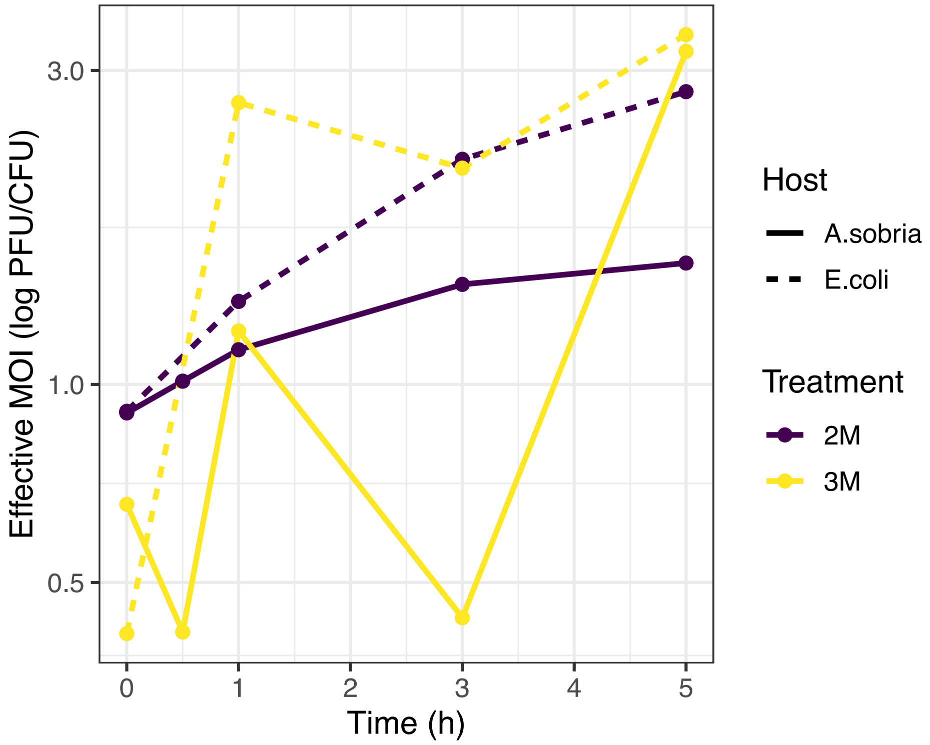


Fig. S4| Effective multiplicity of infection (MOI) dynamics over time in dual predator systems. The log-transformed ratio of PFU to CFU (log PFU/CFU) was used to estimate effective MOI across a 5-hour infection period. Lines represent treatment groups 2M and 3M in each host background (*A. sobria* and *E. coli*), highlighting temporal changes in predator-to-prey ratios and potential differences in infection efficiency and host susceptibility.

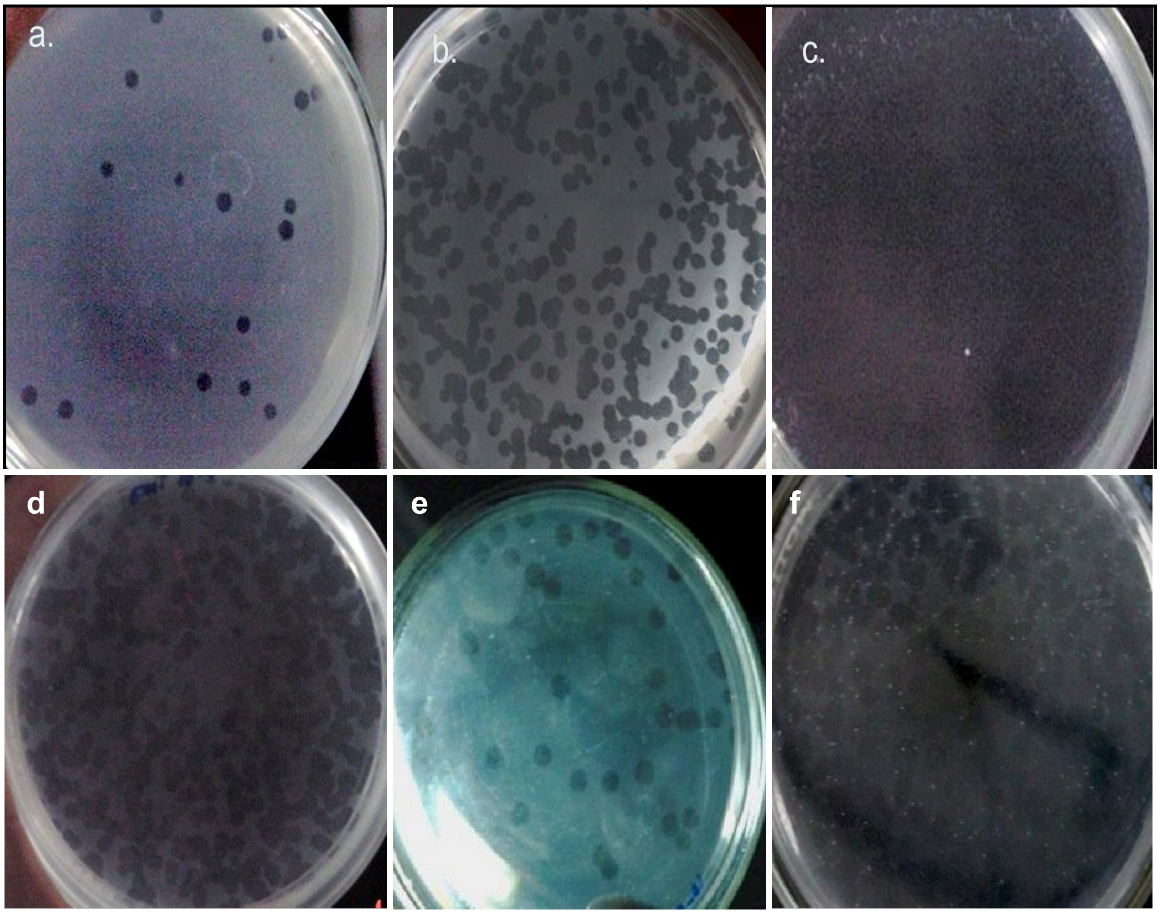


**Fig. S5. Plaque Enumeration**: BALO *EMS* on 2º host *A. sobria* in DLA plates. (a, e) At 4 hr; (b) at 8 hr; (d) at 16 hr; (c, f) at less than 24 hr of incubation, lawn of host bacterium almost completely saturated with plaques.
